# Minimizing variability in the filament middle cerebral artery occlusion model in C57BL/6 mice by surgical optimization - the PURE-MCAo Model

**DOI:** 10.1101/2024.01.22.576769

**Authors:** Sodai Yoshimura, Maximilian Dorok, Uta Mamrak, Antonia Wehn, Eva Krestel, Igor Khalin, Nikolaus Plesnila

## Abstract

**BACKGROUND:** In the intraluminal filament middle cerebral artery occlusion (fMCAo) model, there is considerable variability in infarct volumes, especially in C57BL/6 mice, which often lack the P1 segment of the posterior cerebral artery (PCA) and therefore develop not only MCA but also PCA area infarcts after fMCAo. Another factor contributing to infarct volume variability is collateral flow to the MCA area. The aim of this study was to establish an optimal surgical method to reduce the infarct volume variability in C57BL/6 mice.

**METHODS:** C57BL/6 mice were subjected to 60 min of fMCAo with cerebral blood flow monitored by laser Doppler fluxmetry. The influence of the common carotid artery (CCA) ligation, filament morphology, and the pterygopalatine artery (PPA) ligation on lesion volume and neurological severity score 24 hours after reperfusion were assessed.

**RESULT:** The use of filaments with appropriate length of coating and ligation of the PPA while maintaining perfusion of the CCA prevented the development of infarcts in the PCA area, resulted in pure MCA infarcts (68.3±14.5mm^3^, 26.1±3.6% of the hemisphere with Swanson’s correction) and reduced the variability of infarct volumes by more than half to 13.9% of the standard deviation divided by mean.

**CONCLUSIONS:** Using improved surgical methods with suitable filaments to induce MCA occlusion in mice, we were able to produce <u>P</u>CA area-<u>u</u>naffected <u>re</u>producible infarcts exclusively in the MCA area with reduced variability (PURE-MCAo). Our results may thus help to increase the reproducibility of the fMCAo model and reduce the number of animals required in preclinical stroke research.

## Introduction

The intraluminal filament middle cerebral artery occlusion (fMCAo) model in rodents is widely used for preclinical research of ischemic stroke [1]. This model can reproduce a condition similar to human large vessel occlusion by blocking the blood flow in the MCA from inside the blood vessel. Moreover, by removing the filament, cerebral blood flow can be restored, making the model suitable for studies on reperfusion injury [2, 3]. These features are particularly valuable as they recapitulate the clinical situation of a rapid restoration of blood flow following a period of occlusion achieved by mechanical thrombectomy, a treatment method that has spread widely around the world in recent years [4]. However, the ischemic lesion volume after fMCAo is highly variable, especially in mice [5]. This high variability necessitates the use of a high number of mice to achieve statistically significant differences between experimental groups, or it may lead to false-negative results, consequently hindering the translation of preclinical studies to human treatment.

Among others, one of the major contributing factors to this issue is the significant variation in the development of the P1 segment of the posterior cerebral artery (PCA), which connects the basilar artery to the PCA, across individuals [6]. It is reported that around 50% of C57BL/6 mice, which is the most commonly used strain in preclinical research, lack the ipsilateral P1 segment of the PCA [7]. Since the PCA branches off from the internal carotid artery (ICA) in rodents, infarctions in the PCA area occur in mice lacking the P1 segment of the PCA when the blood flow of the CCA is interrupted during fMCAo surgery, and the CCA is typically ligated or clipped either permanently or temporarily to reduce collateral flow and achieve a more consistent infarction in mice [8]. When an infarct in the MCA area is accompanied by an infarct in the PCA area, the infarct volume is markedly larger compared to an infarct in the MCA area alone [9].

A couple of studies showed that maintaining the blood flow of the CCA during the occlusion period and using a filament with a relatively short length of coating (1 to 2 mm) may prevent infarction in the PCA area, regardless of whether the P1 segment is absent or not [9, 10]. However, it is of concern that maintaining the blood flow in the CCA may lead to an increase in collateral flow to the MCA area [8].

Some time ago, Chen et al. showed that even if the blood flow to the CCA is maintained during the occlusion period, it is possible to reduce the collateral blood flow to the MCA area to a certain extent by blocking the blood flow of the pterygopalatine artery (PPA), which is not usually blocked in mouse fMCAo surgery because of its technical demands [11].

Taking all these into consideration, we hypothesized that the best approach to reduce the variability of infarct volumes after fMCAo would be the combination of maintaining the flow of the CCA during the occlusion period, blocking the flow of the PPA, and using a filament with an appropriate length of coating. To our knowledge, this study is the first to provide a detailed examination of the utility of combining these surgical methods in the fMCAo model in mice. As such, the aim of this study is to verify whether this combined method allows to achieve more consistent infarct volumes compared to conventional surgical methods.

We conducted a comprehensive comparison between our combined fMCAo method and two widespread methods developed by Koizumi [2] and its modified version suggested by Longa [3], the methods using CCA ligation permanently and temporarily, respectively. And we demonstrated the importance of combining two methods by comparing that with the two non-combined methods: specifically, PPA ligation + non-optimal (long tip) filament, no ligation + optimal (short tip) filament.

For a robust evaluation of all five methods, we analyzed infarct volumes, regional cerebral blood flow (rCBF), and neurological severity scores (NSS). Additionally, we explored several surgical techniques to facilitate PPA ligation, a procedure generally considered challenging in mice.

## METHODS

### Animals

Male C57BL/6 mice (8-10 weeks old, 20-26g) were purchased from Charles River Laboratories (Sulzfeld, Germany). Animals were kept in a room with 12h light/12h dark cycle with free access to food and water. All experimental procedures were reviewed and approved by the Animal Ethics Board of the Government of Upper Bavaria.

### Experiment design and surgical methods

In this study, we compared the variability of lesion volumes in our suggested combined method of fMCAo with other methods. Key points of our suggested method are the follows: we do not ligate CCA but ligate PPA and used filaments with short coating tip (1.5mm). Thus, we aimed to stop the blood flow of MCA without affecting the PCA flow and to reduce the collateral blood flow to the MCA area during the occlusion period. A standardized protocol, including anesthesia, rCBF measurements, and duration of the ischemia (60 min), was used in all animals (Fig. 1A). In order to assess influence of surgical interventions on infarct volume and mouse behavior, we performed behavioral tests and Nissl staining of coronal sections 24 hours after reperfusion (Fig. 1A). The difference between the groups was only related to the details of the surgical procedures (Fig. 1B): permanent CCA ligation + short tip filament (Koizumi) group, temporally CCA ligation + short tip filament (Longa) group, PPA ligation + Long tip filament (PL) group, No ligation + Short tip filament (NS) group, and a combination of PPA ligation + short tip filament (PURE-MCAo: PCA-Unaffected REproducible MCAo) group.

**Figure. 1.**
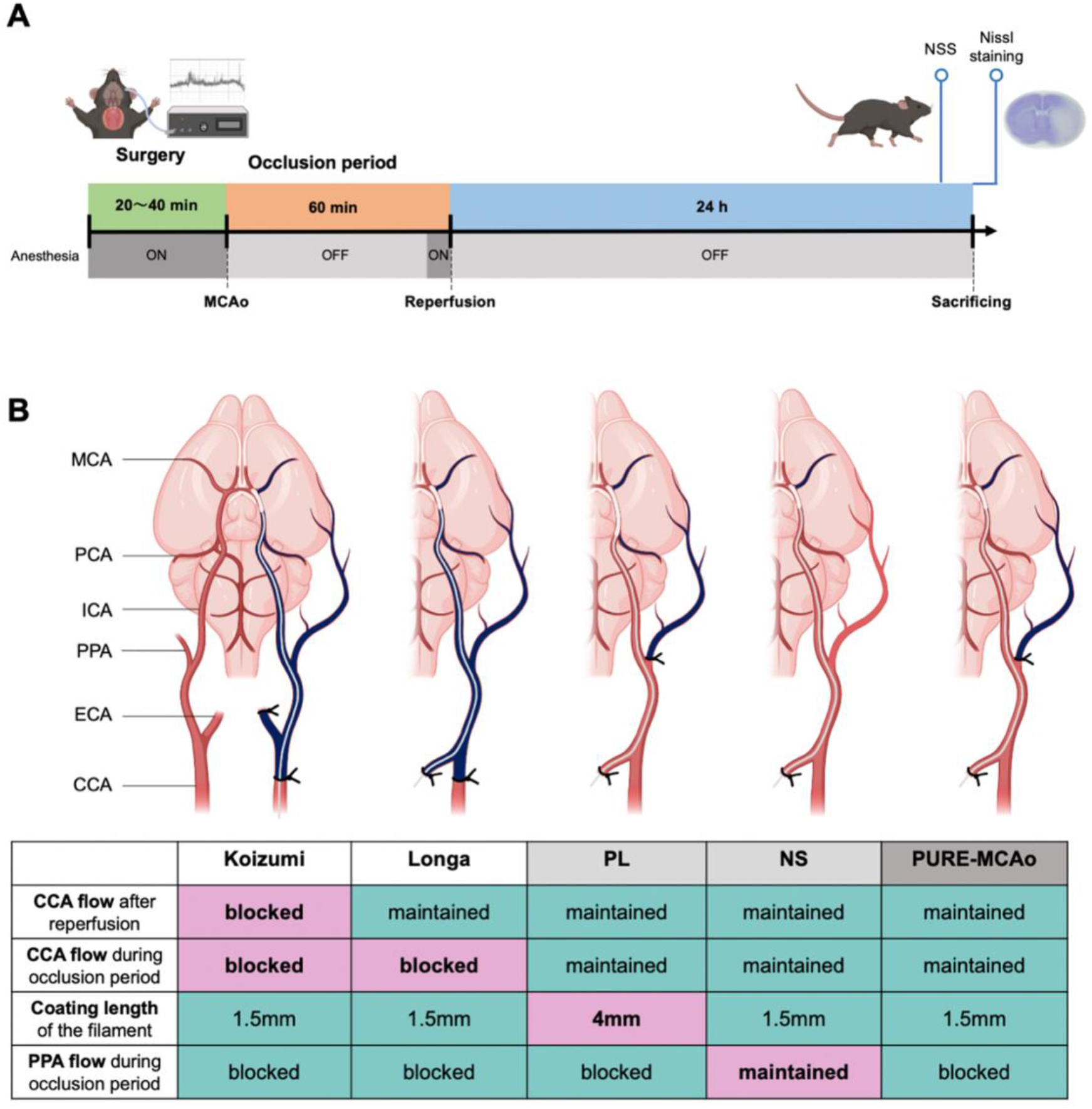
Experimental design and groups description. **A.** Experimental plan: Anesthetized C57BL/6 mice (5 groups × 10 mice) were subjected to 60 minutes of filament middle cerebral artery occlusion (fMCAo). Regional cerebral blood flow (rCBF) was measured during surgery. 24 hours after reperfusion, neurological severity score (NSS) was evaluated, and mice were sacrificed followed by a collection of snap-frozen brain tissue. The coronal brain sections were stained by Nissl staining for further evaluation. **B.** A schematic representation of surgical approaches and respective table with explanations of 5 methods of fMCAo model which were subsequently compared. Namely, Koizumi - permanent common carotid artery (CCA) ligation + short tip filament (1.5 mm); Longa - temporally CCA ligation + short tip filament. Two methods of comparison (without CCA ligation): PL - <u>P</u>PA Ligation + <u>L</u>ong tip filament (4 mm); NS - <u>N</u>o ligation + <u>S</u>hort tip filament. And suggested improved method composed of a combination of PPA ligation and short tip filament - PURE-MCAo (PCA-Unaffected REproducible MCAo).

### Anesthesia

Animals were initially anesthetized with 5% isoflurane and maintained on 2% isoflurane, 30% oxygen, and 70% air. Body temperature was maintained at 36-37°C using a feedback-controlled heating pad (Heater Control Module, FHC, Bowdoinham, ME, USA).

### rCBF Measurement

rCBF was measured using laser Doppler fluxmetry. A skin incision was made from the left ear to the left eye over the skull and the fascia of the temporal muscle was incised. The temporal muscle was then retracted to expose the skull surface. A straight microtip laser Doppler probe (MT B500-0L240, Perimed, Järfälla, Sweden) was attached to the exposed area with a biocompatible adhesive (COSMO CA-500.130, Weiss Chemie + Technik GmbH & Co. KG, Haiger, Germany). rCBF was continuously monitored during the surgery (Perimed Periflux System 5000, Perimed, Järfälla, Sweden). Changes in rCBF during the experimental procedure were normalized to baseline values and reported as a percentage of the initial rCBF. The LabChart software package (v8.1.22, ADInstruments, Sydney, Australia) was used for analysis.

### Filament Middle Cerebral Artery Occlusion

In all animals, the left fMCAo was occluded.

Koizumi group - Animals were secured in a supine position. 1 cm skin incision was made in the middle of the neck. The ECA and CCA were exposed and ligated, and the ICA and its branch, the PPA, were exposed. To later fix the filament, a prepared knot was placed on the CCA, and a vascular clip (Micro Serrefines No,18055-04, Fine Science Tools INC, Vancouver, Canada) was temporarily applied to the proximal ICA. A small incision was made in the CCA, and the filament with 1.5 mm of silicone coating tip (7-0 Max MCAO suture Re L12 PK5, Doccol Corporations, MA, USA) was inserted toward the ICA until a drop in rCBF, indicating that the filament reached the bifurcation of the MCA and the ACA, was confirmed with the laser Doppler flowmetry. The prepared knot was then tightened. The wound was sutured with 6-0 silk thread.

Longa group - After the skin incision, the ECA was exposed, ligated at two locations on the distal side just before the bifurcation of the lingual artery and external maxillary artery, and then cut on the segment between the two ligations and flipped. A prepared knot was placed at the origin of the ECA. The ICA and the PPA were exposed. Blood flow to the flipped ECA was blocked by pulling the prepared knot placed at the origin of the ECA and the suture used to ligate the distal ECA in opposite directions. A small incision was made in the flipped ECA, and the filament with 1.5 mm of silicone coating tip was inserted from the incision toward the ICA until it reached the bifurcation of the MCA and ACA. The CCA was ligated during the occlusion period in this group.

PPA ligation + Long tip filament (PL) group - In this group, while CCA was kept intact, the PPA was temporarily ligated during the occlusion period. The filament with 4 mm of silicone coating tip (7-0 large MCAO suture Re L34 PK5, Doccol Corporations, MA, USA) was used only in this group. Other steps were the same as Longa group.

No ligation + Short tip filament (NS) group - In this group, neither CCA nor PPA were ligated. The filament with 1.5 mm of silicone coating tip was used. Other steps were the same as Longa group.

PPA ligation + short tip filament (PURE-MCAo) group - In this group, while CCA was kept intact, PPA was temporarily ligated during the occlusion period. The filament with 1.5 mm of silicone coating tip was used. Other steps were the same as Longa group.

During the occlusion period, animals were awakened from anesthesia. After 1 hour of occlusion period, animals were anesthetized again and the filament was removed to achieve reperfusion.

Generally, the ligation of the PPA in mice is considered to be highly challenging. However, we found that by retracting the hyoid bone and severing the occipital artery (OA), it is possible to secure sufficient working space, allowing for the safe ligation of the PPA. Technically, by lightly tying the junction of the stylohyoid muscle and the hyoid bone with a suture and retracting the hyoid bone rostrally, the anatomical structures in the vicinity of the skull base, including the PPA, can be better exposed. Additionally, by cutting the OA, more lateral retraction becomes possible, allowing the exposure of the PPA from its origin to the point where it penetrates the skull base. Combining these two techniques makes the ligation of the PPA, which has been considered difficult in mice, relatively easy to achieve while preserving the surrounding anatomical structures. Also, the retraction of the hyoid bone makes it easier to expose the distal side of the ECA, allowing for visualizing the bifurcation of the distal ECA. This ensures that a sufficient length of the ECA can be secured for filament insertion (Fig. 2A-E).

**Figure 2.**
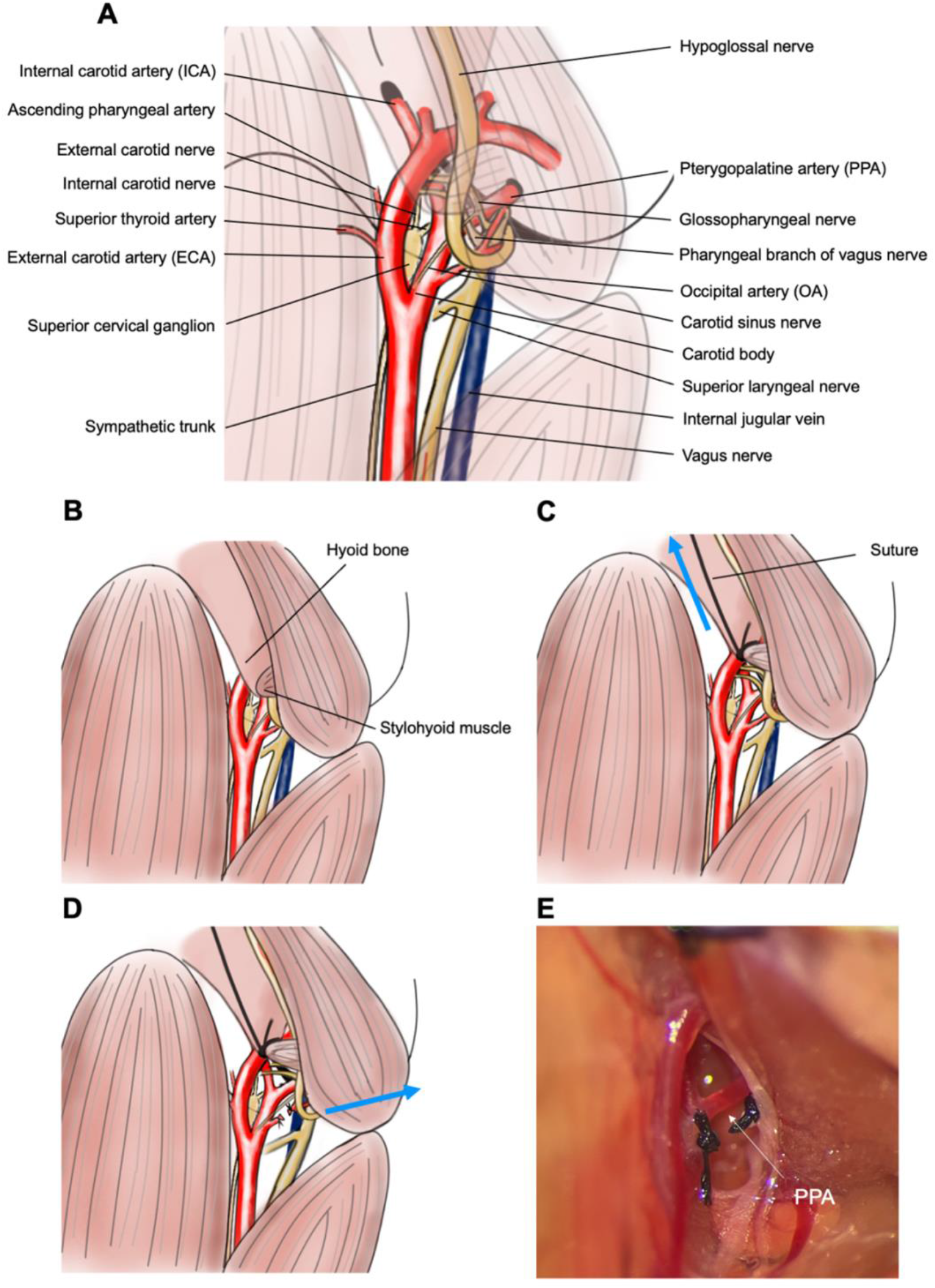
An illustrative description of the surgical procedure to expose the PPA. **A.** Illustration of detailed anatomical exposition of the fMCAo surgical field (drawn by Yoshimura. S.). **B.** The hyoid bone and the posterior belly of the digastric muscle can obstruct the exposure of the PPA. **C.** A suture is inserted between the hyoid bone and the stylohyoid muscle, gently ligating the tip of the hyoid bone and retracting it rostrally, thereby expanding the vertical field of view. **D.** By transecting the OA and retracting the posterior belly of the digastric muscle laterally, the horizontal field of view can be broadened. **E.** The actual surgical field prior to ligation of the PPA.

### Lesion Volume Measurement

Following fMCAo surgery and reperfusion, the brains were harvested 24 hours post-surgery. The extracted brains were immediately snap-frozen in dry ice and stored at -20°C. Brain sections were prepared using a cryostat (Cryostar NX70, Thermo Fisher Scientific, MA, USA), obtaining 12 slices per brain with a thickness of 10 μm and an interval of 750 μm, starting 2.8 mm anterior to Bregma. The sections were stained with Nissl staining [1], and images were captured using a light microscope (ZEISS Axio Imager M2, Carl Zeiss, Jena, Germany) with an 1× objective (EC Plan-Neofluar 1×/0.025 M27, Carl Zeiss, Jena, Germany). The infarct area was quantified using the AxioVision Rel 4.8 software (Carl Zeiss, Jena, Germany). The infarct volume (mm³) was calculated by multiplying the sum of the infarct areas from each slice by 0.75 mm. To account for the effects of brain edema, the corrected infarct volume (%) was also determined using the Swanson’s correction method [12]. ImageJ (version 1.53t, National Institute of Health, Bethesda, MD, USA) was employed to create averaged images from multiple photographs of Nissl-stained brain slices. 3D reconstruction of the averaged images was done using IMARIS software (Oxford Instruments, MA, USA).

### Neurological Severity Score (NSS)

Neurological function was assessed in all mice with the NSS 24 hours after reperfusion. We modified the Bederson score[13] as described below to allow for video-based assessment, facilitating scoring by third parties.

### 0: No neurological deficit

1. Slight difference in strength between the limbs on the side contralateral to the lesion, with asymmetrical gait during walking.

2. Inability to walk straight due to contralateral paralysis caused by the lesion, resulting in a circling movement.

3. Unable to move from the spot, rotating in place.

4. Severe contralateral paralysis caused by the lesion, unable to maintain posture.

When the degree of neurological symptoms was determined to be positioned between each score, respective scores of 0.5, 1.5, 2.5, 3.5 were assigned.

### Selective intra-arterial injection

A catheter (LDPE-Schlauch für die Medizintechnik, K28640 inner diameter: 0.28mm, outer diameter: 0.61mm, Reichelt Chemietechnik GmbH & Co, Heidelberg, Germany) was seared and stretched to narrow the tip. For PCA injection, the filament with 1.5mm of coating tip was placed at the bifurcation of the ACA and MCA, using the same method as fMCAo surgery of the PURE-MCAo group. Subsequently, the catheter was inserted from the ECA and its tip was positioned in the ICA, after which it was secured with a suture over the blood vessel. For the PPA injection, the catheter was inserted from the ECA and its tip was placed in the PPA. The ICA and the CCA were ligated to prevent the injection into areas outside the target region due to backflow. Following deep sedation with intraperitoneal injection of a solution containing medetomidine (1.0 mg/kg), fentanyl (0.1 mg/kg), and midazolam (10 mg/kg) (MMF), the chest cavity was opened, exposing the heart. After the right atrium was incised and exsanguination euthanasia performed, the ascending aorta was severed. Subsequently, Lycopersicon Esculentum (Tomato) Lectin (LEL, TL), DyLight™ 649 (Vector Laboratories, CA, USA) was injected arterially, at a volume of 150µl, followed by an injection of 150µl of saline via the catheter. The brain was removed and then fixed by immersing in 4% of paraformaldehyde (PFA) for 48 hours. It was then embedded in 4% agarose gel (VWR, PA, USA), and 60 µm sections were prepared using a vibratome (VT1200S, Leica Biosystems, Wetzlar, Germany). These sections were examined with a confocal microscope (ZEISS LSM 980, Carl Zeiss, Jena, Germany). The imaging conditions were set to a 10x magnification (Plan-Apochromat 10x/0.45 M27) with an image matrix of 512 × 512 pixels and a pixel scaling of 1.66 × 1.66 µm. Image depth was adjusted to 8 bits. Images were captured in z-stacks with tile scans, maintaining a slice distance of 10 µm and a total range of 60µm. The laser wavelength was 639nm and detection wavelength was between 641 and 694 nm.

### Statistical Analysis

Statistical analyses were conducted using GraphPad Prism (version 9.4.1, GraphPad software Inc, USA). For comparisons between two independent groups, a Student’s t-test was applied when the data followed a normal distribution, while the Mann-Whitney U test was used for data not following a normal distribution. For comparisons among three or more groups, a one-way Analysis of Variance (ANOVA) was performed for data following a normal distribution, whereas the Kruskal-Wallis H test was used for data not following a normal distribution. As post-hoc multiple comparisons, Tukey’s test or Dunn’s test was performed.

## Results

### The PPA plays an important role for the perfusion of the MCA territory

To investigate whether the PPA pays a role for the perfusion of the MCA territory we measured rCBF during MCA occlusion by laser Doppler fluxmetry in all experimental groups (**Fig. 3A**). Except in the NS group, occlusion of the MCA together with ligation of either the CCA (Koizumi, Longa) or the PPA (PL, PURE-MCAo) resulted in a reduction of rCBF by more than 80%, that is to ischemic levels (**Fig. 3B**). In the NS group (no CCA or PCA ligation) rCBF decreased significantly less than in the other groups (36.3 ± 14.2%, p<0.0001; **Fig. 3B**). Interestingly, rCBF dropped only then to ischemic levels when the PPA was pinched with a forceps (**Fig. 3C**). This experiment suggests that the PPA contributes to MCA territory perfusion. To investigate whether the PPA indeed perfuses the MCA territory, we inserted a catheter into the PPA and injected fluorescent lectin to stain all vessels distal of the injection site (**Fig. 3D**). The strongest fluorescent signal was observed in the meninges, however, also vessels within cerebral cortex were stained (**Fig. 3E**). Additionally, also vessels from the base of the brain heading towards the hypothalamus and the internal capsule showed lectin fluorescence (**Fig. 3F**). Altogether, these data suggest that the PPA plays an important role for the perfusion of cortical and striatal areas traditionally believed to be solely perfused by the MCA or by the anterior choroidal artery, respectively. These results suggest that the PPA may significantly contribute to the variability of the filament MCAo model.

**Figure 3.**
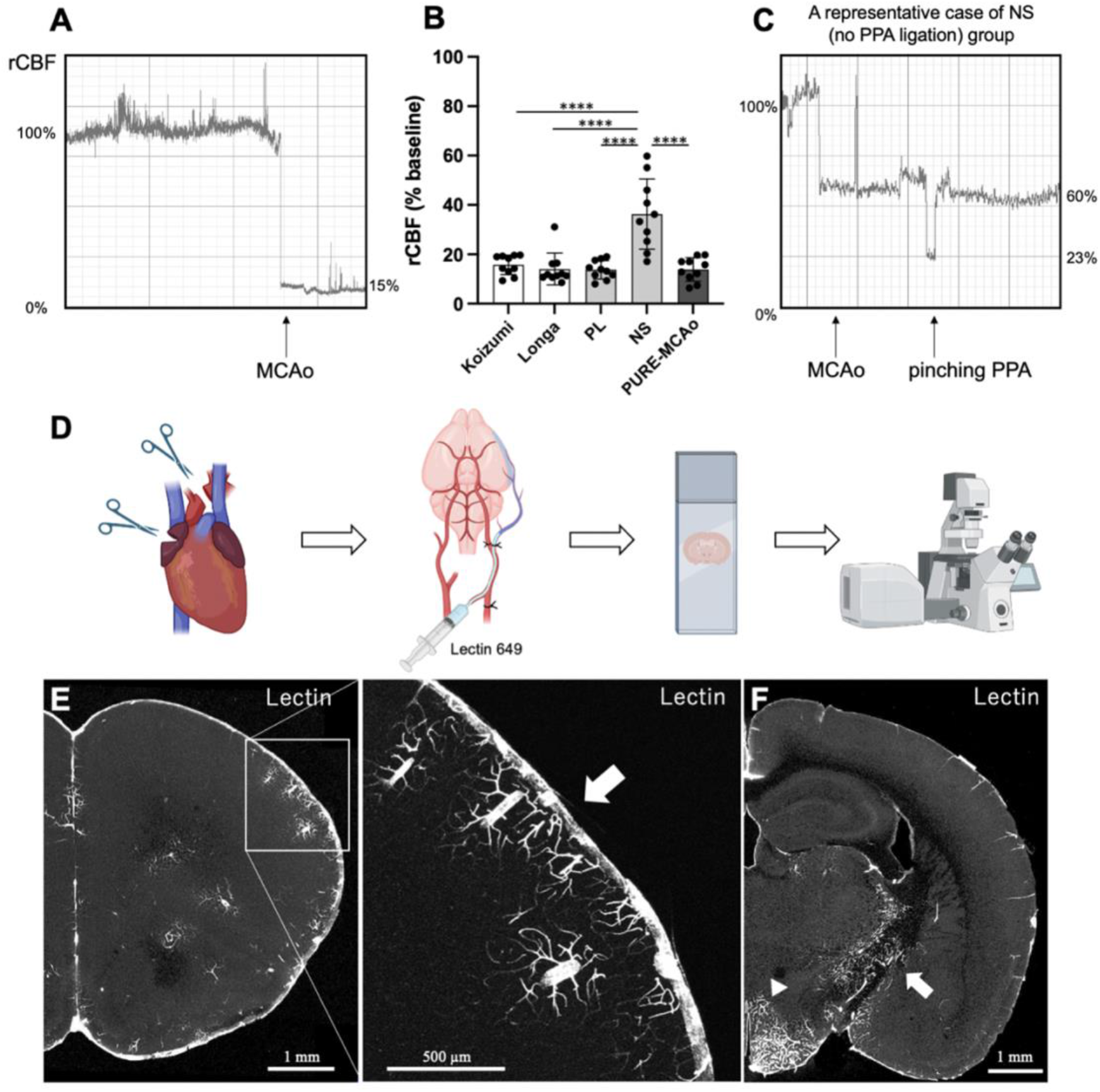
Cerebral blood flow during surgery and selective arterial injection of fluorescent lectin to PPA. **A**. Representative waveform of laser Doppler fluxmetry. Sudden decrease of rCBF is observed right after MCA occlusion. **B.** Quantitative analysis of rCBF during occlusion period. Data are presented as individual values and mean ± SD, 10 mice per group. Statistical analysis was performed using GraphPad Prism^®^, employing one-way ANOVA with a Tukey post-hoc test for multiple comparisons. **** indicates p<0.0001. **C.** A representative case of NS group. After filament MCAo, additional decrease of rCBF was observed when pinching PPA. **D.** Experimental design for identifying brain areas perfused by the PPA. Catheter was introduced into PPA in deeply anesthetized mouse. After the right atrium was incised and exsanguination euthanasia performed, the ascending aorta was severed in order to avoid labelling the rest of the brain vasculature. Subsequently, Lectin 649 was injected via the catheter. Then mouse was fixed by paraformaldehyde (PFA) and collected brain was cut in vibratome. Coronal sections were imaged using fluorescent confocal microscope. **E.** A representative coronal brain section of a naïve mouse injected with lectin. White arrow - signal from meninges. Meninges and cortical vessels were stained. **F.** Vessels within the hypothalamus (arrowhead) and the internal capsule (white arrow) were also stained by lectin injected through the PPA.

### PURE-MCAo exhibits the least variability in infarct volume

As a next step we evaluated the infarct volume following all five MCA occlusion approaches using Nissl staining of coronal brain sections 24 h post-reperfusion. First, we examined the spatial distribution of the lesion (**Fig. 4A**). We observed that the Koizumi, the Longa and the group with long tip filament (PL) all resulted in infarcts including the PCA area, while the NS and the PURE-MCAo group did not show infarctions within the PCA territory (**Fig. 4B**). In the Koizumi, the Longa, and the PL group, PCA infarcts were observed in 6, 3, and 6 out of 10 cases (magenta), respectively. In the NS and PURE-MCAo group not a single mouse suffered a PCA infarct. Accordingly, the infarct volumes in the Koizumi (42.55 ± 11.43%) and the PL group (44.30 ± 11.69%) were significantly larger than in the NS (22.03 ± 7.11%) and the PURE-MCAo group (26.05 ± 3.62%; **Fig.4 D**). These results demonstrate that the number of animals with infarctions within the PCA territory significantly increases the average infarct volume. Additionally, comparing the standard deviation (SD) relative to the mean among all groups, the PURE-MCAo group exhibited the lowest variability (13.88%), which was approximately 50% smaller than that of any of the other groups (Koizumi: 26.87%, Longa: 38.44%, PL: 26.37%, NS: 32.26%) (**Fig. 4E**). Altogether, the PURE-MCAo method induces selective infarctions of the MCA territory with a very low variability.

**Figure 4.**
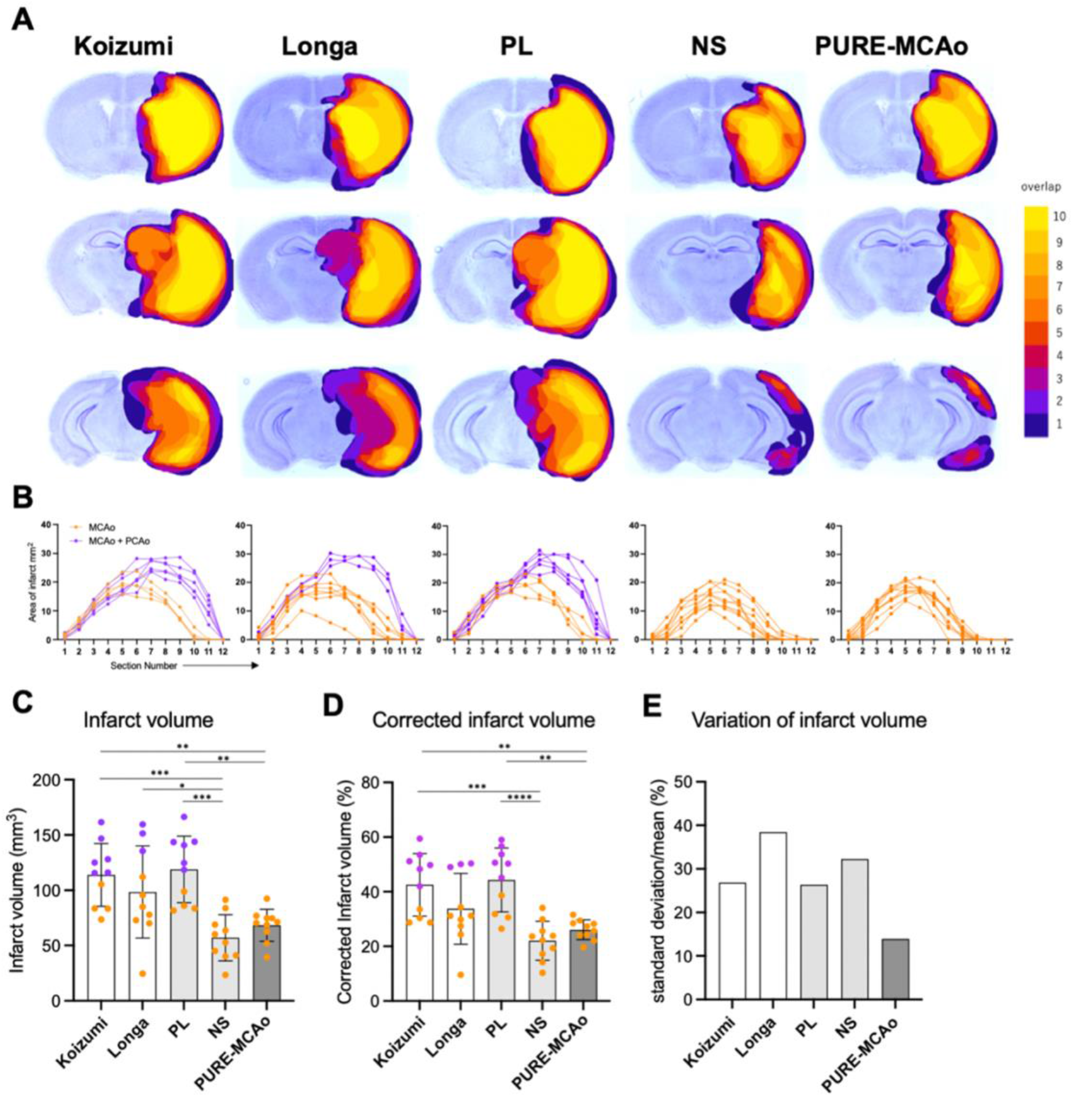
Variability of infarct volume and distribution. **A.** Three representative slices (out of 12 total) from the brain 24h post fMCAo, stained with Nissl staining, are shown from each experimental group (upper: -0.2mm, middle: 1.7mm, lower: 3.2mm from bregma). The color code represents the spatial distribution of lesion overlap, from 1 (dark blue) to 10 (yellow). **B.** Graphical representation of the lesion area at every of 12 brain slices from each animal. Groups corresponds to A. Orange: MCA area infarct, magenta: MCA + PCA area infarct. **C,D.** Quantitative analysis of absolute infarct volume (C) and corrected to edema infarct volume (D) among all experimental groups. Data are presented as individual values and mean ± SD, 10 mice per group. Statistical analysis was performed using GraphPad Prism^®^, employing one-way ANOVA with a Tukey post-hoc test for multiple comparisons. *: p<0.05, **: p<0.01, ***: p<0.001, ****: p<0.0001. Color code – orange: animals with infarct at MCA area, magenta: animals with infarct at MCA + PCA area. **E.** Comparison of variation of infarct volume (standard deviation/mean) in each group.

### The PURE-MCAo method markedly decreases the number of animals required in experiments

Using the G*Power software, we calculated the required sample size for comparative experiments on infarct volume between two groups using each method. For the settings, an α value of 0.05 (P < 0.05) and a power of 0.8 were used. With the conventional Koizumi method and Longa method, to detect a 20% difference, 20 and 39 mice/group would be needed, respectively. In contrast, the PURE-MCAo method would require only 6 mice/group. Even when aiming to detect differences of 30-50%, the PURE-MCAo method would reduce the group size by at least 50% compared to the traditional models. The newly developed PURE-MCAo method, regardless of the magnitude of the effect detected, may therefore very effectively reduce the number of experimental animals needed to detect a robust difference between experimental groups in pharmacological or mechanistic studies, which are traditionally underpowered due to large date scattering (**Fig. S1**).

### Neurological severity score, survival rate, and surgery time

Comparing the neurological outcome of all experimental groups 24 hours post-reperfusion, we found that a higher neurological severity score (NSS) was present in the groups with MCA + PCA territory infarcts (**Fig. 5 A**), namely, Koizumi, Longa, and PL. The lowest NSS (1.1 ± 0.84) was observed in the NS group, which was statistically different from the Koizumi (p = 0.0191) and the PL group (p = 0.0069). That a high NNS was caused by a combination of MCA and PCA infarctions was also reflected by the finding that the proportion of animals with PCA + MCA area infarct per group showed a high correlation with a worse NSS score (r = 0.945; p = 0.015; data not shown). Further, those techniques which caused MCA and PCA infarcts had the largest variability in neurological outcome (NSS grades 1-4). The NP group which showed only MCA infarctions had also a large variability in terms of outcome (NSS grades 0.5-2.5) most likely due to insufficient reduction of blood flow during MCAo. The smallest variability in neurological function was observed in the PURE-MCAo group, the only group with selective MCA infarctions and reductions of cerebral blood flow below the ischemic threshold during MCAo.

**Figure 5.**
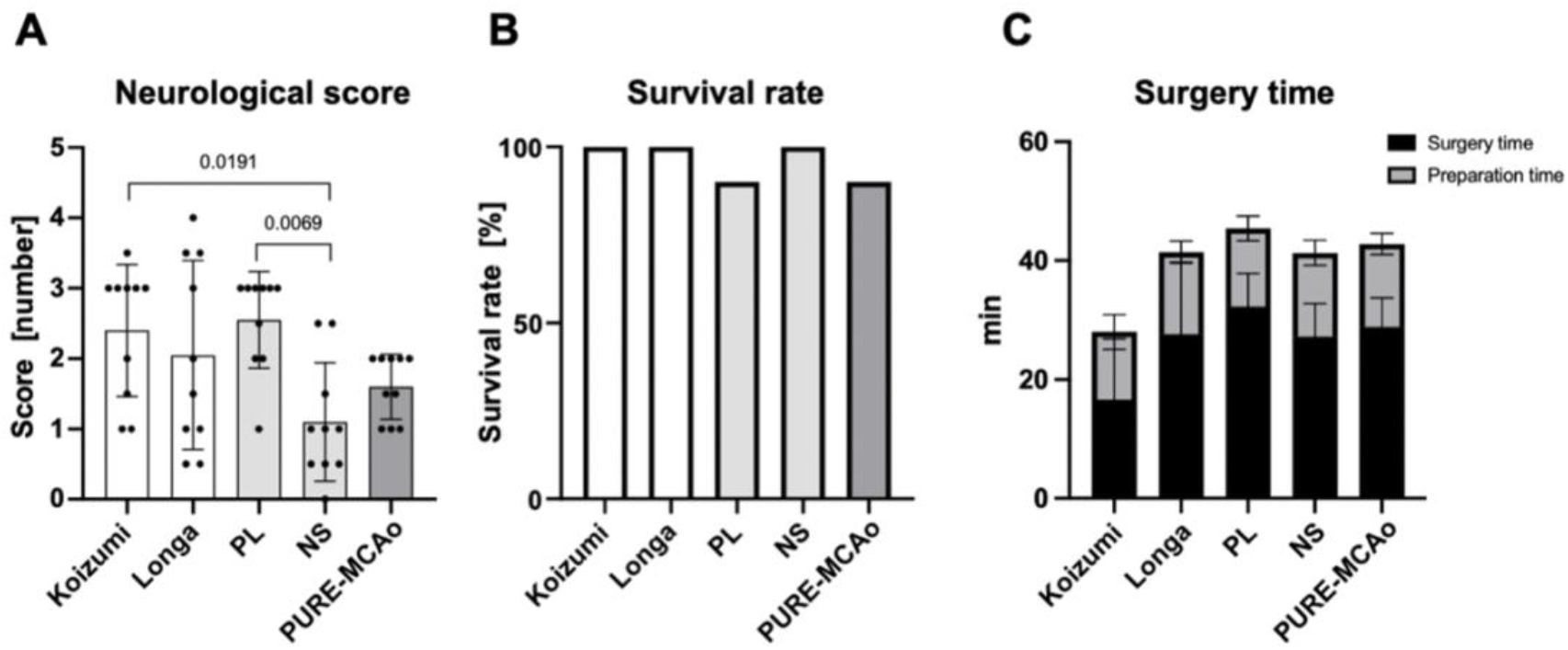
Neurological severity score and other parameters. **A.** A graph presenting quantitative analysis of neurological severity score (NSS) evaluated 24 hours post-reperfusion. Data are presented as individual values and mean ± SD, 10 mice per group. Statistical analysis was performed using GraphPad Prism®, employing one-way ANOVA with a Tukey post-hoc test for multiple comparisons. **B.** A graph presenting proportional presentation of survival rate within 24 hours following reperfusion. **C.** A graph presenting surgery and preparation time spent for model performance. Data are presented as mean ± SD, 10 mice per group.

Overall the rate of survival 24 hours after MCAo was 90% or higher in all groups. One animal died in PURE-MCAo group within 24 hours of reperfusion due to subarachnoid hemorrhage and one animal died in PL group in the second half of the observation time most likely due to intracranial hypertension. In the other three groups, all mice survived for 24 hours after reperfusion (**Fig. 5B**).

The Koizumi method allowed to occlude the MCA after only 16.6±10.2 min, while all other methods to occlude the MCA took much longer, i.e. between 27.2±5.6 and 32.3±5.5 min. (**Fig. 5C**). Nevertheless, the longer anesthesia times had no adverse effects on infarct volumes or neurological function.

### Comparison of MCA vs. MCA + PCA area infarction

Next, we sought to understand which functional areas of the mouse brain where affected by MCA and MCA+PCA infarcts and to what extent additional PCA infarcts influenced animal behavior. For this purpose, we analyzed data from all 50 mice regardless of the surgery technique used for MCAo and subdivided these animals into two groups: mice with MCA infarcts (n=35) and mice with MCA + PCA infarcts (n=15; **Fig. 6 A**). The mean infarct volume in the MCA group was 71.1 ± 21.2 mm^3^, while in the MCA + PCA group the infarct volume was almost twice as large (138.5 ± 16.8 mm^3^; p<0.0001; **Fig. 6B**). Also the NSS was significantly different between these two groups: the NSS score was more than one unit higher in the MCA+PCA group (1.6 ± 0.9 in the MCA infarct group vs. 2.8 ± 0.79 for the MCA + PCA infarct group; p<0.0001; **Fig. 6C**). These data clearly indicate that incidental PCA occlusions massively worsen the histopathological and behavioral outcome following filament MCAo in mice.

**Figure 6.**
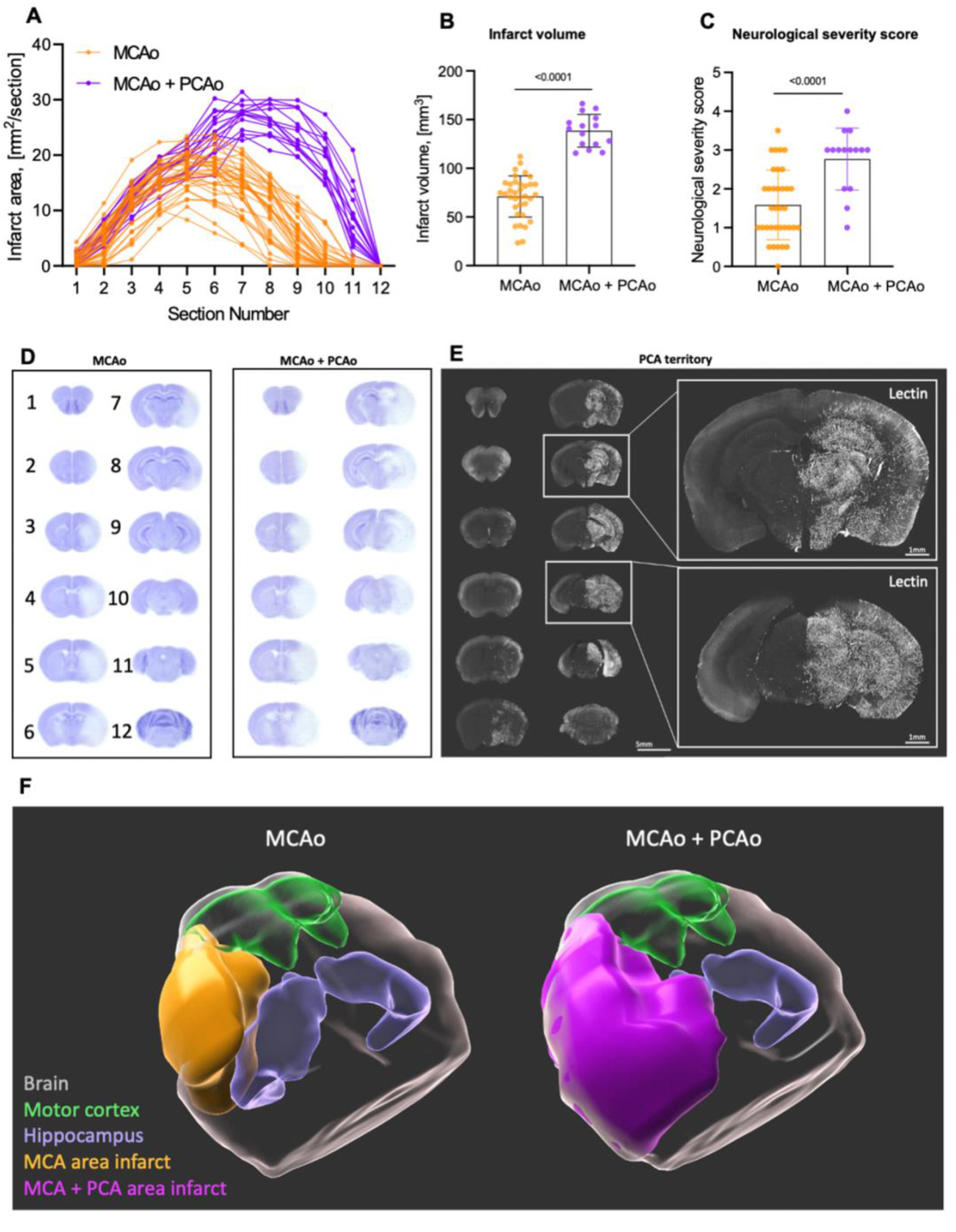
Comparative analysis of posterior middle cerebral artery (PCA) infarction, behavior, and territory. **A.** Graphical representation of the lesion area at every of 12 brain slices from each animal. Orange - MCA area infarct group, magenta - MCA + PCA area infarct group. **B.** Quantitative analysis of infarct volume of MCA area infarct *vs.* MCA + PCA area infarct. **C.** A graph presenting quantitative analysis of neurological severity score. Data are presented as individual values and mean ± SD. **B, C.** - Data are presented as individual values and mean ± SD, MCA area infarct - 35 mice, MCA + PCA area infarct – 15 mice. Statistical analysis was conducted using an unpaired t-test in GraphPad Prism®. **D.** Averaged images of 12 Nissl stained coronal sections from 35 mice with MCA area infarct (left) and 15 mice with MCA+PCA area infarct (right). **E.** A representative confocal image of coronal section of respective to Lectin injection hemisphere. Lectin was injected via the internal carotid artery after filament MCA occlusion with subsequent brain fixation and imaging as was depicted at Fig. 2D. This enabled directing tracer exclusively via PCA area. **F.** 3D reconstructed images of the brains, based on the averaged Nissl-stained slices shown in Figure D. The areas of infarction are color-coded as follows: MCA area infarct (orange), MCA + PCA area infarct (magenta), motor cortex (green), and hippocampus (blue).

To investigate the brain regions affected by MCAo or MCAo+PCAo, average images of the MCA and MCA + PCA infarct groups were generated and compared (**Fig. 6D**). We found no infarcts in the two most rostral (1) and dorsal sections (12) proving that the whole infarct area was covered by the currently used sectioning protocol. We found no difference between groups in the four subsequent rostral sections (2 to 5). However, there were large differences between groups in slices 7 to 11. Most importantly, we observed only cortical and striatal infarcts in the MCA group at slices 6 to 10, while in the MCA+PCA group, the hippocampus and the thalamus were also damaged.

To directly assess the PCA territory in mice, we performed selective MCAo using the PURE-MCAo technique and injected fluorescent lectin via the internal carotid artery (**Fig. 6E**). Our data showed that the PCA supplies blood flow to the posterior third of the cerebral cortex, hippocampus, midbrain, thalamus, and hypothalamus. This fully matches with the infarcted brain areas following MCAo+PCAo assessed by Nissl staining. For better visualization of the brain areas affected by MCAo or MCAo-PCAo we performed 3D reconstructions of the 2D data shown in Fig. 6D and added the motor cortex and the hippocampus based on the Allan Brain atlas (**Fig. 6F**). These data confirm that the infarcts caused by MCAo+PCAo are much larger than those induced by MCAo alone and affect areas not perfused by the MCA, i.e. the hippocampus. An interesting subsidiary finding of this analysis is that the motor cortex was only minimally affected in both groups. Hence, it may not be purposeful to use motor function as a primary functional outcome parameter after MCAo.

## Discussion

The filament MCA occlusion model was initially developed by Koizumi et al. in rats [2]. In this model, permanent CCA ligation is required as the filament is inserted from the CCA. Reperfusion is then achieved through collateral circulation via the anterior communicating artery and P1 segment of PCA. Subsequently, Longa et al. [3] introduced a method that avoids sacrificing the CCA by inserting the filament from the ECA. While this method allows for a more complete reperfusion due to anterograde flow from the CCA, studies comparing the two methods have reported no significant differences in infarct volume or survival rates [14]. Furthermore, the Longa technique has longer surgery and anesthesia times and was therefore avoided by some researchers [14]. Later, with the widespread use of genetically modified mice, the fMCAo model has also been widely adopted in mouse studies. However, the difference in size between rats and mice has led to more variations in surgical techniques. One of the biggest differences in the technique to induce MCAo between rats and mice is the ligation of the PPA. While this procedure is commonly performed in rats to reduce collateral blood flow to the brain, it is considered difficult and is therefore generally not conducted in mice. Instead, CCA ligation during the occlusion period is widely performed not only in the Koizumi method but also in the Longa method in mice [8]. As a result, frequent infarctions in the PCA region have been observed in C57BL/6 mice, which often lack the P1 segment of the PCA [6]. Consequently, when using the Koizumi or Longa method in mice, a combination of MCAo and PCAo will occur. This issue is often disregarded and in most investigations MCA and PCA territory infarctions are summed up and reported to be MCA infarcts.

In our current study, we quantified the volumes of MCA and MCA+PCA infarcts and revealed that occlusions of the MCA+PCA result in infarct volumes approximately twice as large as those observed after specific MCAo. Our results also show that when using the most widely used techniques to occlude the MCA in mice based on the Koizumi and Longa techniques, about 50% of mice suffer unintended PCA infarctions. Consequently, the standard deviation of the infarcts induced with these techniques has a large variability of 40% or more. Large infarcts, which affect also neighboring vascular territories, result is missing collateral perfusion of the dorsal MCA territory and may thus compromise therapeutic interventions addressing mechanisms improving collateral blood flow. Large group sizes to overcome the unnecessary high variability of the currently used MCAo models pose ethical questions, waste resources, and delay drug development programs. Hence, the use of the improved technique to induce MCAo, PURE-MCAo, may significantly improve the current situation and result in more and more reliable and reproducible data.

Furthermore, our study demonstrates that PCA infarctions significantly affect neurological outcome following MCAo, suggesting that not only infarct size but also neurological scores may yield inconsistent results when the PCA is affected. Since PCAo in the Koizumi and Longa models occurs by chance, false positive or false negative results may also occur purely by chance. This hinders data consistency and serves as a major obstacle in translating preclinical findings to clinical applications.

In our view, the current study clearly indicates that mice with PCA infarcts following MCAo need to be excluded from analysis, particularly when testing pharmacological compounds. When using conventional fMCAo methods in C57BL/6 mice, however, this would result in the exclusion of about 50% of all operated animals. Even if animals can be excluded immediately or soon after surgery by measuring CBF with laser speckle imaging or assessing the ischemic territory by MRI [9][15], this is an ethically and economically impractical solution.

The results of the current study suggest that a relatively simple modification of the surgical technique used to induce MCAo in mice, termed PURE-MCAo, may offer a fundamental solution to the above outlined problem and its wide-reaching consequences for translational stroke research. Our approach combines the findings of Akamatsu et al.[9], who reported on the importance of maintaining CCA flow during the MCA occlusion period without CCA ligation and using a filament with an appropriate coating length, and Chen et al.[11], who demonstrated that PPA ligation reduces collateral flow to the brain. In this study, we confirmed that maintaining CCA blood flow during the MCA occlusion period is a prerequisite to avoid PCA infarctions. Regardless of whether CCA ligation was permanent or temporary to the occlusion period, the Koizumi and Longa groups, in which CCA ligation was performed, both showed PCA infarctions. Furthermore, our examination of the PL group demonstrated that even when the flow of the CCA is maintained, PCA territory infarctions cannot be avoided unless the filament has the appropriate length of coating. Lastly, by comparing the NS and the PURE-MCAo group, we confirmed that PPA ligation contributes to the decrease of rCBF and the variability of the infarct volumes in the cerebral cortex. Only the combination of these three elements we were able to achieve consistent MCA territory ischemia with minimal variability concerning infarct volumes and functional outcome.

Our current study also provides novel results regarding the perfusion of the mouse brain. While it had been previously demonstrated that the PPA provides collateral blood flow to the MCA territory by angiography [16], the specific regions of the brain supplied by the PPA have not been investigated in detail. To examine which areas of the mouse brain are perfused by the PPA, we performed intraarterial fluorescent lectin injection into PPA and demonstrated that mice have functional anastomosis between the meninges and the cerebral cortex. We also demonstrated that the PPA provided collateral blood flow to the hypothalamus and the internal capsule.

Another interesting finding of our study is that MCAo in mice does in most cases not affect the motor cortex responsible for limb movement, regardless of the presence or absence of PCA infarction. This suggests that impairments of limb movements observed after fMCAo cannot be caused by damage to the motor cortex, but most likely to more downstream structures like white matter fibers of the pyramidal tract including the internal capsule. It is known that the internal capsule is mainly perfused by perforating branches from the ICA and the anterior choroidal artery. Hence, the worse neurological function in mice with MCAo + PCAo are well explained by occlusion of these arteries. On the other hand, our observation that the PPA also supplies blood to the internal capsule may explain why animals in the PURE-MCAo group exhibited higher NSS scores with less variability as compared to the NS group, where the PPA was not occluded.

As acknowledged by experienced surgeons, details of a surgical procedure may significantly alter outcome. In the current study we developed a surgical procedure which allows the ligation of PPA in mice, a procedure considered to be technically very challenging, with ease. We discovered that by retracting the hyoid bone rostrally and cutting the occipital artery to allow retraction toward the outside, the PPA can be sufficiently exposed, enabling a relatively safe ligation. Insufficient exposure during PPA ligation or clipping attempts can pose risks of damaging surrounding structures such as the lingual branch of the glossopharyngeal nerve, the recurrent laryngeal nerve, or the carotid sinus nerve, emphasizing the importance of proper exposure.

## Conclusion

Current filament MCA occlusion models in mice cause infarctions in the MCA and PCA territories thereby producing highly unreliable results with large variations. The newly developed PURE-MCAo model induces selective MCA occlusions with high reliability and reduces the variability of infarct volumes by approximately 50%. Thus, the PURE-MCAo has the potential to eliminate most of the shortcomings of current fMCAo models, to reduce animal numbers, and to significantly increase the reproducibility and reliability in the field of experimental stroke research.

## Abbreviation

ANOVA: Analysis of Variance
CCA: common carotid artery
ECA: external carotid artery
fMCAo: filament middle cerebral artery occlusion
ICA: internal carotid artery
MCA: middle cerebral artery
OA: occipital artery
PCA: posterior cerebral artery
PFA: paraformaldehyde
PPA: pterygopalatine artery
rCBF: regional cerebral blood flow
SD: standard deviation

## Acknowledgement

This work was supported by the Deutsche Forschungsgemeinschaft (DFG, German Research Foundation) grant 457586042.

## Conflict of interest

Authors declare no conflict of interest

---

**Figure S1.**
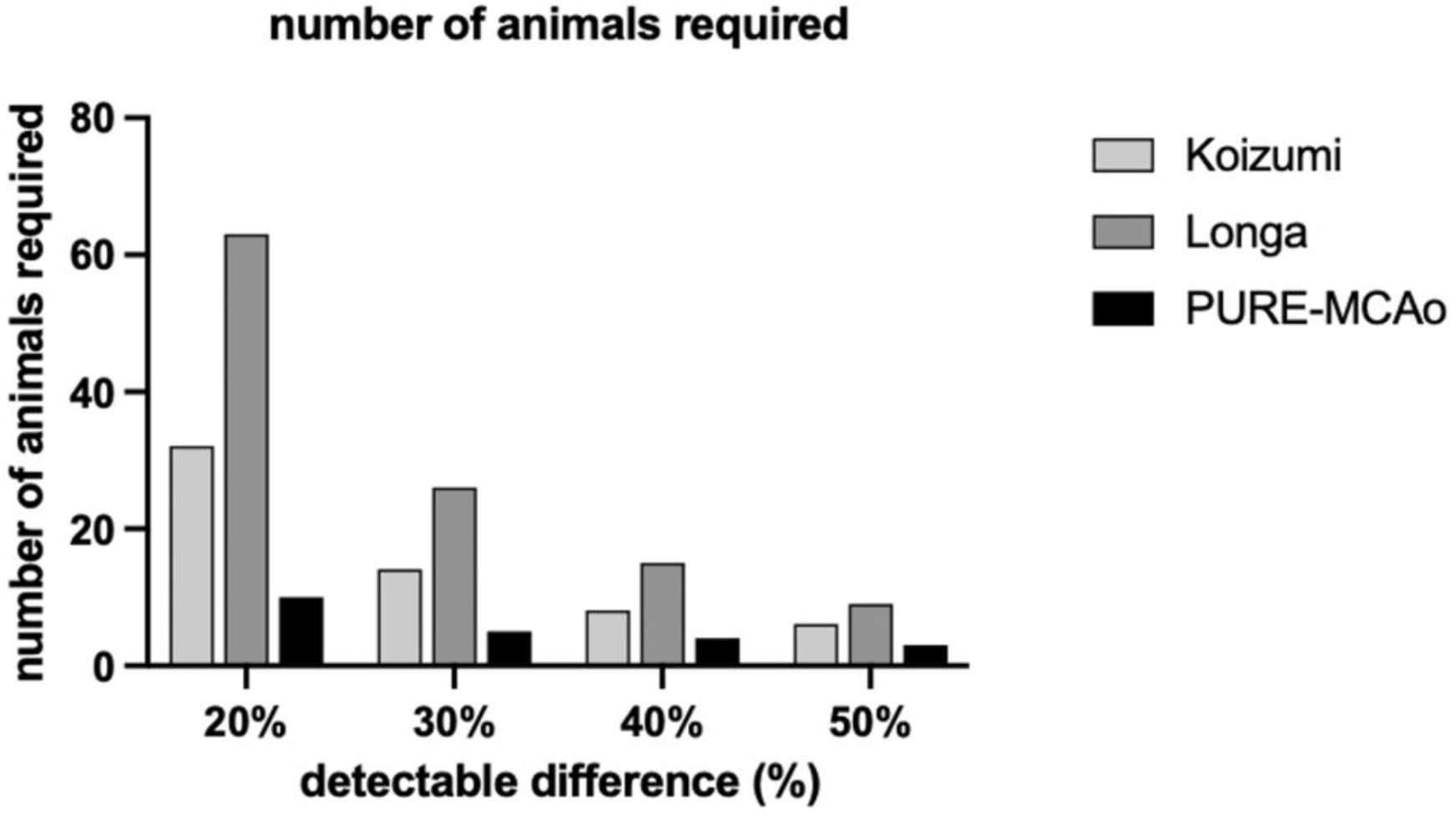
Comparison of animal numbers required in experiments across methods. A graph shows the number of animals required to achieve each detectable difference (20%, 30%, 40%, and 50%) in infarct volume among the Koizumi method, Longa method, and PURE-MCAo method. Sample size calculations were conducted using the G*Power software, with settings of α = 0.05 and power = 0.8.

## Notes

### Competing Interest Statement

The authors have declared no competing interest.

